# Previously unknown regulatory role of extracellular RNA on bacterial directional migration

**DOI:** 10.1101/2024.07.11.603110

**Authors:** Victor Tetz, Kristina Kardava, Maria Vecherkovskaya, Alireza Khodadadi-Jamayran, Aristotelis Tsirigos, George Tetz

## Abstract

Bacterial directional migration plays a significant role in bacterial adaptation. However, the regulation of this process, particularly in young biofilms, remains unclear.

Here, we demonstrated the critical role of extracellular RNA as part of the Universal Receptive System in bacterial directional migration using a multidisciplinary approach, including bacterial culture, biochemistry, and genetics.

We found that the destruction or inactivation of extracellular RNA with RNase or RNA-specific antibodies in the presence of the chemoattractant triggered the formation of bacterial “runner cells» in what we call a “panic state” capable of directional migration. These cells quickly migrated even on the surface of 1.5% agar and formed evolved colonies that were transcriptionally and biochemically different from the ancestral cells. We have also shown that cell-free DNA from blood plasma can act as a potent bacterial chemoattractant. Our data revealed a previously unknown role of bacterial extracellular RNA in the regulation of bacterial migration and have shown that its destruction or inhibition triggered the directional migration of developing and mature biofilms towards the chemoattractant.

## INTRODUCTION

Bacterial migration is central for all aspects of bacterial life, allowing bacteria respond to various physical, chemical, biological, and mechanical stimuli. It is a key element that regulates bacterial behaviors such as virulence, biofilm formation, and colonization of new areas (^1^). The primary way for bacteria to migrate is through the activation of swimming, swarming, twitching, gliding, and sliding motility (^2–4^). Both bacteria within biofilms and free-floating bacteria can migrate; however, regulation of migration varies between biofilm-growing and planktonic microorganisms (^5^).

Biofilms are the dominant mode of bacterial life in nature, playing multiple roles in microbiota maintenance and protection of bacteria from antibiotic and immune assault (^6–10^). Directed cell migration from biofilms is a key process involved in host colonization and the systemic spread of infection; thus, understanding factors that regulate this process is of vital importance (^11,12^).

For directional migration, biofilm-growing bacteria must not only sense the chemoattractant but also escape from their biofilm by a complicated process known as biofilm dispersal (^10,13,14^). First, a liquefaction of extracellular polymeric substances (EPSs), in which bacteria within biofilms are embedded, must occur (^15^). Next, even motile bacteria are in a non-motile state within biofilms; therefore, in order for such cells to migrate, the transition between the sessile and motile lifestyle should be activated (^5,12^). However, the transition is complicated due to biofilm multicellular organization and the high regulation rate of bacteria.

One way to coordinate the switch from a biofilm lifestyle to dispersal is via quorum sensing, a communication strategy for cells within biofilms that involves chemical signaling(10). Quorum sensing is the predominant regulator of mature biofilm dispersal, which is cell density-dependent and acts through the inhibition of matrix compound synthesis, increased production of surfactants, and EPS degradation (^16,17^).

Another well-accepted key regulator of the biofilm-to-motility transition is the secondary messenger cyclic di-guanosine-monophosphate (c-di-GMP)(^14^). Many studies have demonstrated that a high intracellular c-di-GMP concentration promotes biofilm formation by enhancing the production of EPSs, which restrict bacterial motility (^18,19^). Conversely, lower intracellular c-di-GMP concentrations promote directional migration and biofilm migration and dispersal by enhancing cell motility by upregulating expression of flagellar protein synthesis and flagellum assembly (^20–23^). The c-di-GMP pathway is an important way by which bacteria respond to diverse external stimuli; however, the resulting interbacterial cascades are much better understood than the actual factors that decrease c-di-GMP’s intracellular concentration (^24^).

Although unrelated to the regulation of directional cell motility, another way to disperse biofilms and regulate bacterial migration is through modulation of the extracellular matrix and extracellular DNA in particular (^25–27^). The importance of extracellular nucleic acids for bacterial biofilms is although not being under the debate, but various research suggests different functionalities ranging from helping to form sticky polymers required for bacterial adhesion, being used in horizontal gene transfer, or even having receptive and regulatory roles (^13,28–31^). Considering the role of polymeric cell-free DNA as a mechanical component of the extracellular matrix, different articles have described the use of DNA-degrading nucleases as a strategy to facilitate biofilm dispersal (^30,32^).

The factors that lead to transition between the sessile and motile lifestyles of bacteria within microbial biofilms are insufficiently studied, particularly for young and immature biofilms for which activation of directional migration cannot be explained with quorum sensing and regulation of EPS viscosity. Therefore, the discovery of novel regulatory systems that control bacterial directional migration is extremely important.

Until recently, extracellular RNA was not known to be associated with regulation of bacterial motility or directional migration. However, we recently reported the unique role of a special type of extracellular RNA that controls major bacterial characteristics and governs the response to a diverse array of environmental, chemical, and physical factors (^28^). We identified that at least some part of extracellular nucleic acids possesses a receptive and regulatory function, and we have demonstrated the existence of nucleic-acid-based receptors (Teazeled receptors, TezR) which are part of the Universal Receptive System (^28,33^). Here, we investigated the effect of the inactivation of extracellular RNA on the regulation of directional migration, particularly in the context of biofilm dispersal, using *Bacillus pumilus*, a species with peritrichous flagella capable of different straight or curved runs (^34^).

## RESULTS

### Inactivation of extracellular RNA triggers bacterial migration and biofilm resettlement toward a chemoattractant

We studied the effect of RNase on *Bacillus pumilus* VT1200 migration toward a human plasma used as a chemoattractant. First, we used a custom-made four-compartment Petri dish model to study bacterial migration. The agar in each of the three experimental compartments was individually supplemented with human plasma, RNase, or both RNase and human plasma, while the control compartment contained 0.9% sodium solution. *B. pumilus* suspension was inoculated in the center of the Petri dish, and in these settings, bacteria demonstrated a high selectivity of migration (Fig. 1). The only zone into which bacteria migrated was the zone containing both human plasma and RNase. In our previously conducted studies, we found that the addition of RNase to the agar did not result in RNase penetration into *B. pumilus (*^28^).

**Figure 1:**
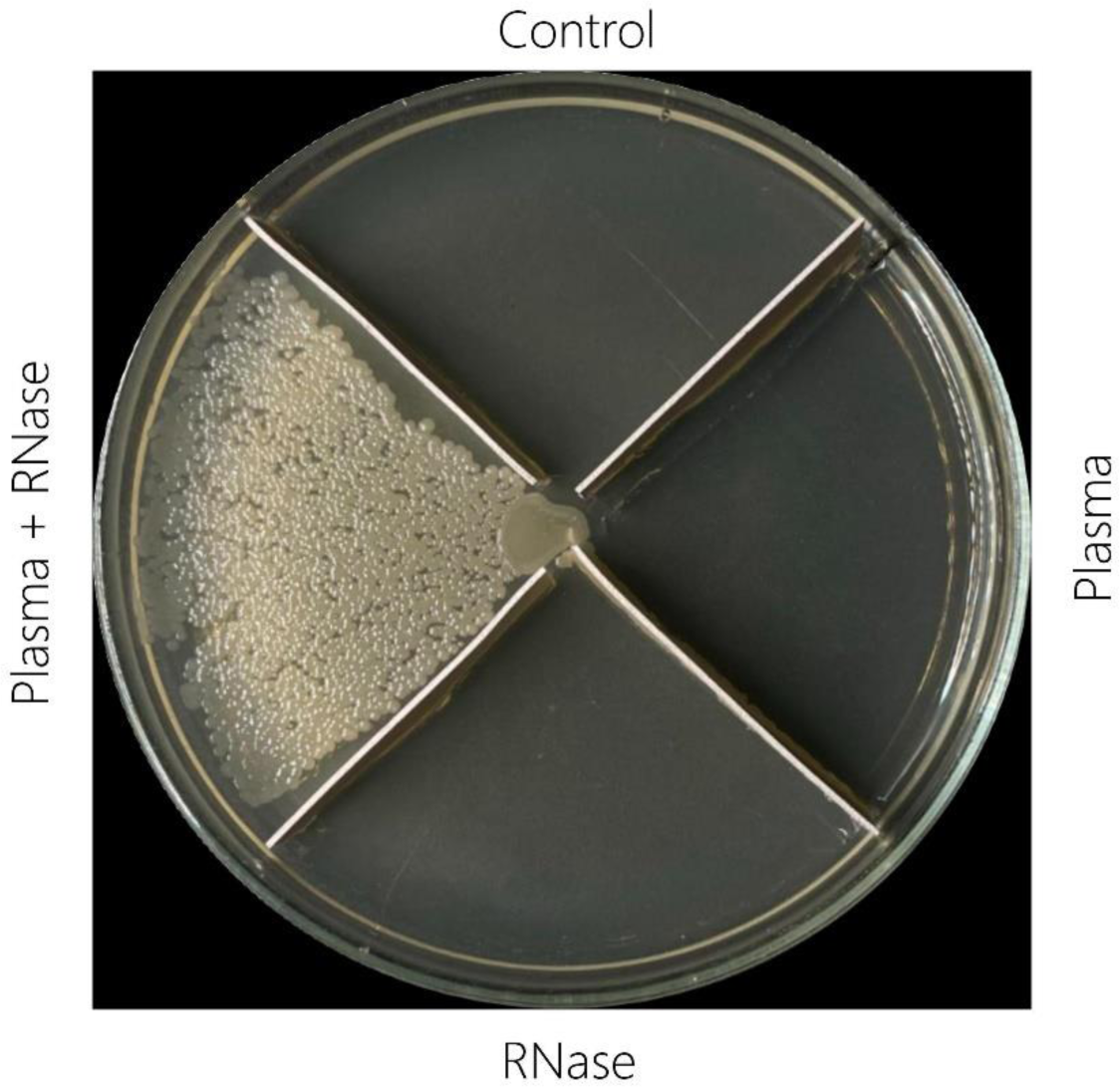
The effect of RNase addition on *B. pumilus* migration toward agar supplemented with plasma used as a chemoattractant. Bacteria were inoculated in the center of a four-compartment Petri dish, and the compartments were filled 24 h later with agar containing a 0.9% aqueous solution of NaCl (control), plasma (plasma), **(3)** a 0.9% aqueous solution of NaCl with RNase 50.0 µg/mL (RNase), or **(4)** plasma and RNase 50.0 µg/mL (Plasma + RNase). Pictures are from representative experiments, which were performed in triplicate.

We next showed that the destruction of extracellular RNA with RNase triggers the surface-motility induction of biofilms of different ages. *B. pumilus* biofilms were formed on agar supplemented or not supplemented with RNase for 24 h and 48 h, after which plasma as a chemoattractant was added on the top of 1/3 sector of the Petri dish.

We named ancestral biofilms grown on RNase-free agar with plasma “ancestral RNase^free^ biofilm,” *Bacillus* grown on RNase-supplemented agar with plasma “ancestral RNase^supp^ biofilm,” and evolved colonies grown on RNase-supplemented agar with plasma “evolved colonies.”

Ancestral RNase^free^ biofilms did not show directional migration after adding plasma (Fig. 2A). Under the same conditions, ancestral RNase^supp^ 24- or 48-hour-old biofilms dispersed, forming multiple evolved colonies that grew solely within the zone where plasma was added. In order to exclude the possibility that biofilm directional migration resulted from greater swimming motility due to decreased EPS viscosity due to loss of extracellular RNA, we also added DNase to the agar aimed to destroy extracellular DNA and further decrease biofilm viscosity. However, bacteria cultivated on agar supplemented with both RNase and DNase did not disperse, meaning that alteration of EPS viscosity did not cause resettlement of ancestral RNase^supp^, which echoed with our previous findings that combined use of DNase and RNase might have a opposite regulatory effects on bacterial cells (Supplementary Figure 1) (^35^). To ensure that the observed effects were related to the specificity of the effect of RNA inactivation on biofilm directional migration and resettlement, we conducted the same experiment, but instead of destruction of plasma extracellular RNA with RNase, we inactivated it with anti-RNA antibodies (Fig. 2B). We observed the formation of the evolved clones within a zone supplemented with anti-RNA antibodies and plasma, which confirmed that the inactivation of bacterial extracellular RNA in the medium was a critical element for directed migration.

**Figure 2:**
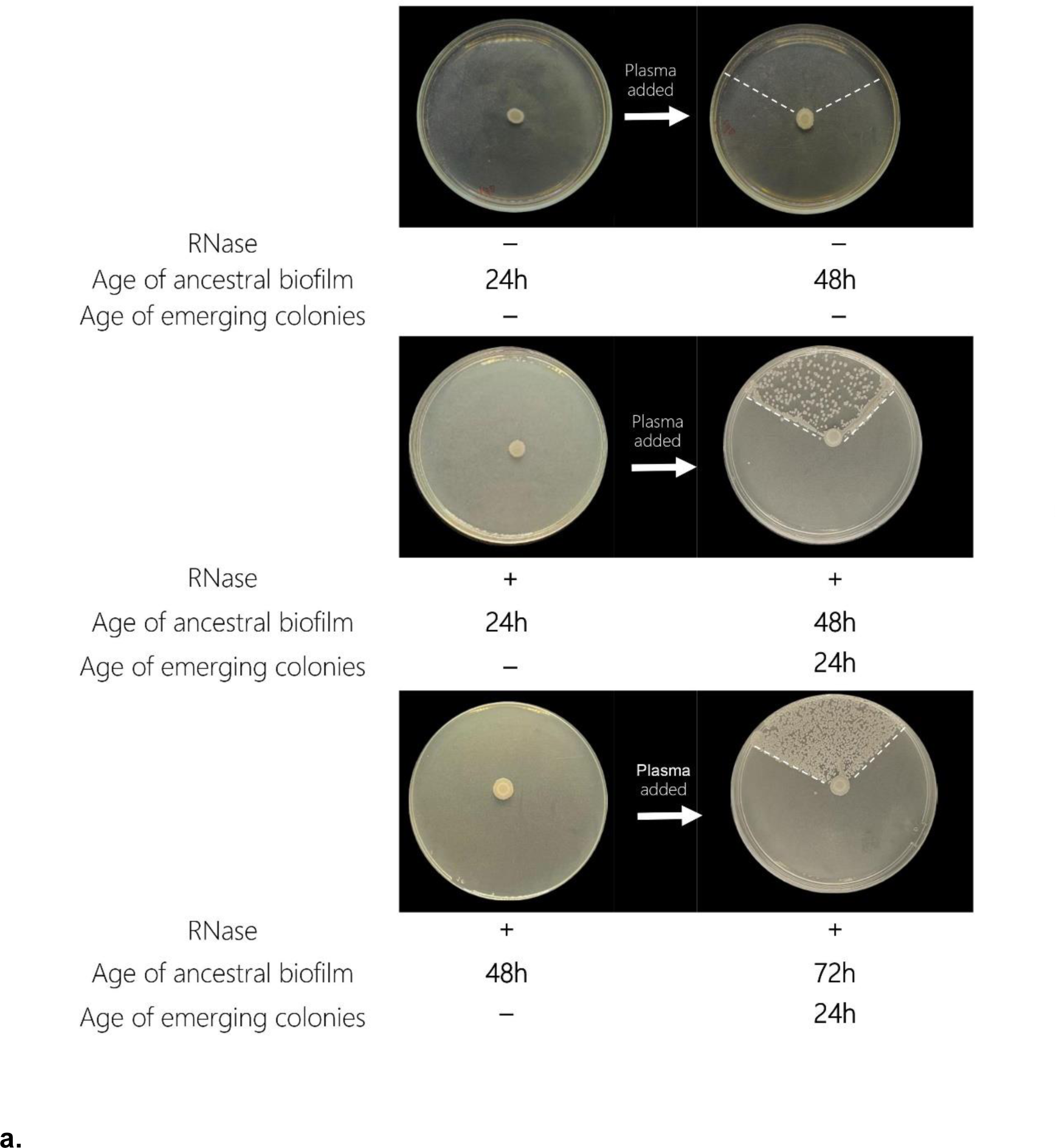
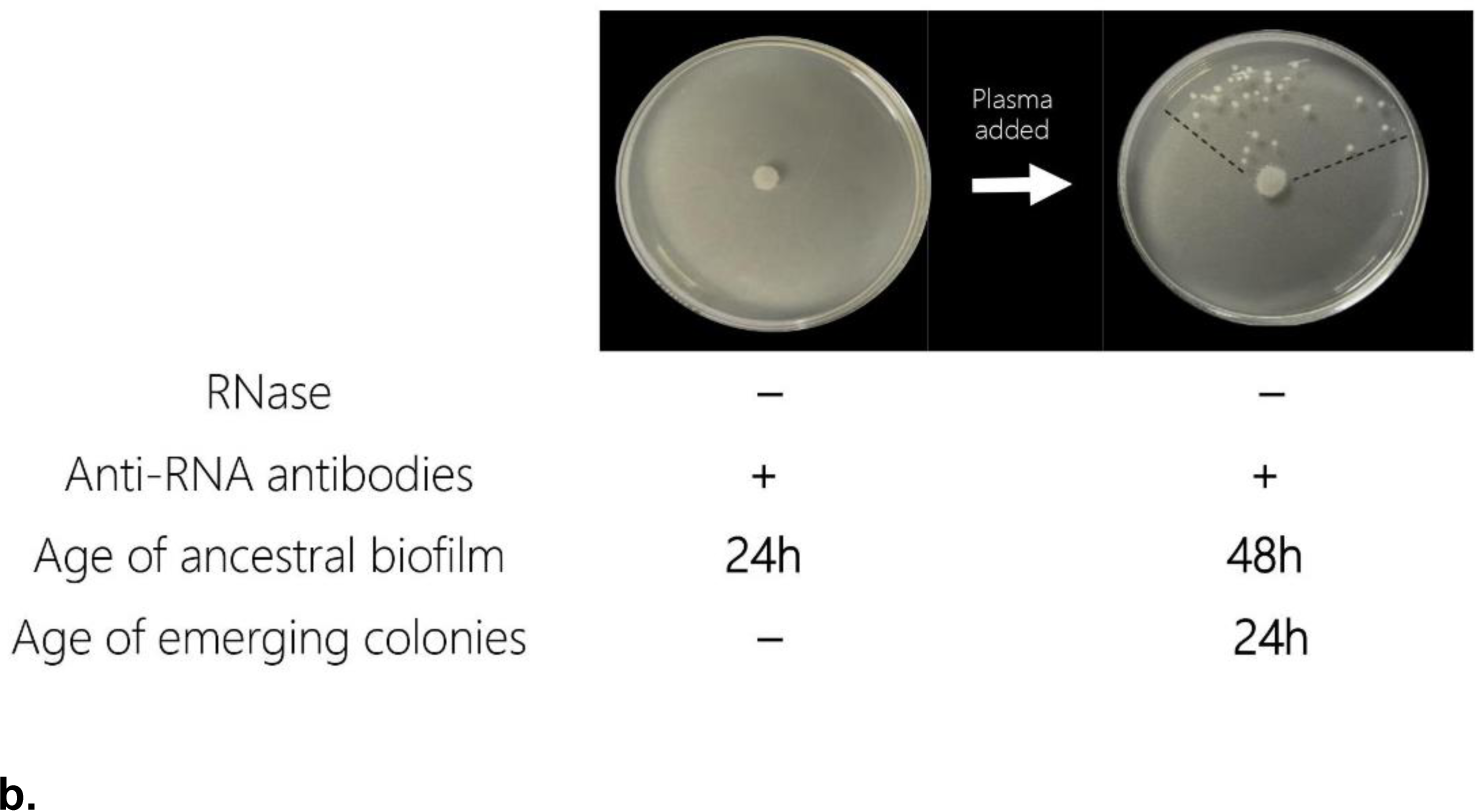
The effect of extracellular RNA destruction and inactivation on biofilm resettlement towards plasma used as a chemoattractant on 1.5% agar. (A) Effect of plasma addition to the agar of different age ancestral RNase^free^ biofilms (AC RNase^free^) or ancestral RNase^supp^ biofilms (AC RNase^supp^) on evolved colony (EC) formation. White dotted lines depict the zone within which plasma was added. (B) Biofilm directional migration after inactivation of extracellular RNA by anti-RNA antibodies. Pictures are from representative experiments, which were performed in triplicate. Dotted lines mark the zone where plasma was added.

### Addition of RNase allows bacterial cell migration over long distances

Next, we studied the distance that bacteria could migrate to a chemoattractant on media supplemented with RNase. We added additional barriers on the Petri dish to increase the maximum distance available for the cells from the initial inoculum to move. Plasma was added to the agar in the form of a path to guide the growth of bacteria from the initial inoculum within the corridor (Fig. 3A). By 24 h, we observed that the cells were dispersing through the entire route, strictly following the chemoattractant, with evolved colonies growing in the zone containing plasma from the original inoculum. The total distance they migrated was 35 cm. This result highlights that the addition of RNase to agar in the presence of a chemoattractant enabled cells to migrate untypically long distances, particularly on the solid 1.5% agar. To study the maximum distance of cell migration, we next inoculated bacteria in a 24-cm-long plate with a total travelable distance of 66 cm and containing RNase-supplemented agar and plasma as a chemoattractant added in the form of a curved path. We found that by 24 h, cells had migrated the whole distance from the initial inoculum, forming microcolonies throughout the entire 66-cm route of the added chemoattractant (Fig. 3B).

**Figure 3:**
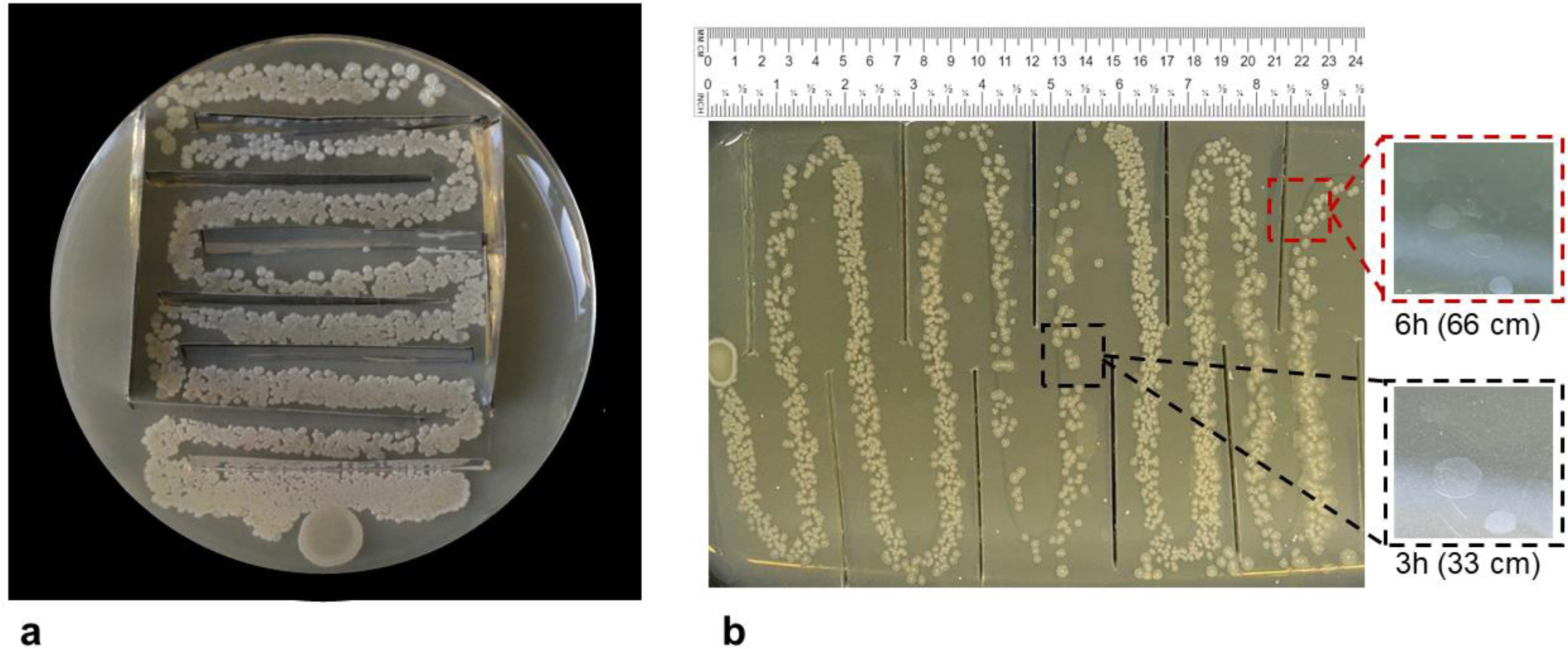
Migration distance of bacteria in RNase-supplemented agar with blood plasma as a chemoattractant. Plasma was added to the agar surface in the form of a curved path using an inoculation loop, guiding the growth of bacteria from the original inoculum along the chemoattractant pathway within the corridor created by plastic inserts placed in the agar. (A) Bacterial growth in a Petri dish that was separated by barriers to increase the possible travelable distance to 33 cm. (B) Bacterial migration in a 24-cm-long plate with barriers that increase the total distance up to 66 cm after 24 h of growth (main image). Black dotted lines mark the magnified zone with the earliest emergence of the most distantly evolved colonies after 3 h of growth (within 33 cm away from the inoculum). Red dotted lines mark the magnified zone with the earliest emergence of the most distantly evolved colonies after 6 h of growth (66 cm away from the inoculum). Pictures are from representative experiments, which were performed in triplicate.

We next studied the time that evolved colonies required to pass half of the distance (33 cm) or the whole distance (66 cm). The appearance of evolved colonies on the agar surface was monitored hourly with a microscope. The earliest visible bacterial growth within 33 cm of the ancestral colony was seen at 3 h, and bacterial growth that had reached the end of the plate was observed after 6 h (66 cm away from the initial inoculum). *B. pumilus* migration reached the edge of the 66 cm plate; however, we believe that this is not the limit of its migratory capability. The migration of *B. pumilus* was also captured over 12 hours of incubation, showing the appearance of distantly evolved colonies (Supplemental Video 1).

### Effects of RNase on sporulation and morphological characteristics of cells

Next, we analyzed what other bacterial characteristics are alerted in cells following extracellular RNA destruction.

Since dispersed cells travel different distances before forming evolved colonies, we compared the characteristics of 24-h-old evolved colonies that had traveled three different distances from the ancestral RNase^supp^ biofilm and compared them to each other and to those of ancestral RNase^supp^ biofilms and ancestral RNase^free^ biofilms (Fig. 4, Supplementary Figure 2).

**Figure 4:**
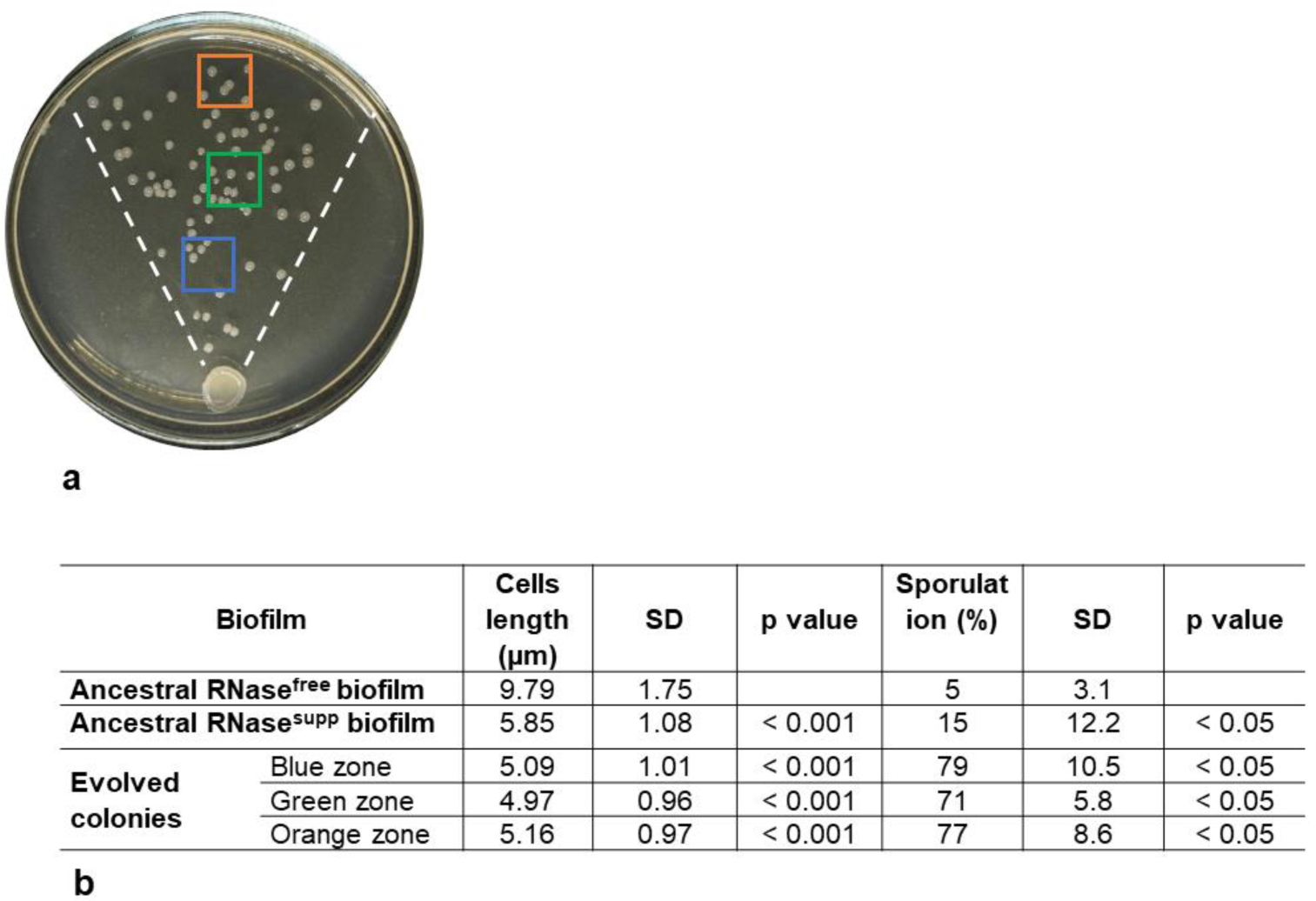
Characteristics of bacteria from 24-h-old ancestral RNase^free^ biofilms, ancestral RNase^supp^ biofilms, and evolved colonies. (A) Evolved colonies at 2 cm, 4 cm, and 6 cm from ancestral RNase^supp^ biofilm are highlighted with corresponding blue, green, and orange rectangles (B) Comparison of bacterial characteristics. Dotted lines mark the zone where plasma was added. Pictures is a representative image from the study performed in triplicate.

When compared to each other, the size of evolved colonies from any of three distances from the ancestral RNase^supp^ biofilm were identical. We found that cultivation of bacteria on RNase-supplemented agar resulted in significantly smaller cell size of bacteria from ancestral RNase^supp^ biofilms and evolved colonies when compared to those from ancestral RNase^free^ (p < 0.05) biofilms. Moreover, bacteria from evolved colonies demonstrated higher sporulation when compared to those from ancestral RNase^supp^ and ancestral RNas^free^ biofilms, regardless of how far they migrated from the ancestral colony.

Similar trends were obtained for 12-h-old biofilms, meaning that the alterations of cell characteristics following extracellular RNA destruction came early and were persistent (Supplementary Figure 3).

### Transcriptome profiling underlying biofilm directional migration

To gain further insight into the regulatory pathways that govern directional migration triggered by bacterial cultivation on agar supplemented with RNase, we compared RNA-seq analyses of *B. pumilus* ancestral RNase^free^ biofilms, ancestral RNase^supp^ biofilms, and evolved colonies. Hierarchical clustering of RNA-seq datasets highlights the largest pairwise Euclidean distance between the ancestral RNase^free^ biofilm and evolved colonies (Fig. 5a). The same trend is clearly evident after multivariate analysis by principal component analysis (PCA), which confirmed the greatest amount of variance between ancestral RNase^free^ biofilm and evolved colonies (Fig. 5b).

**Fig. 5.**
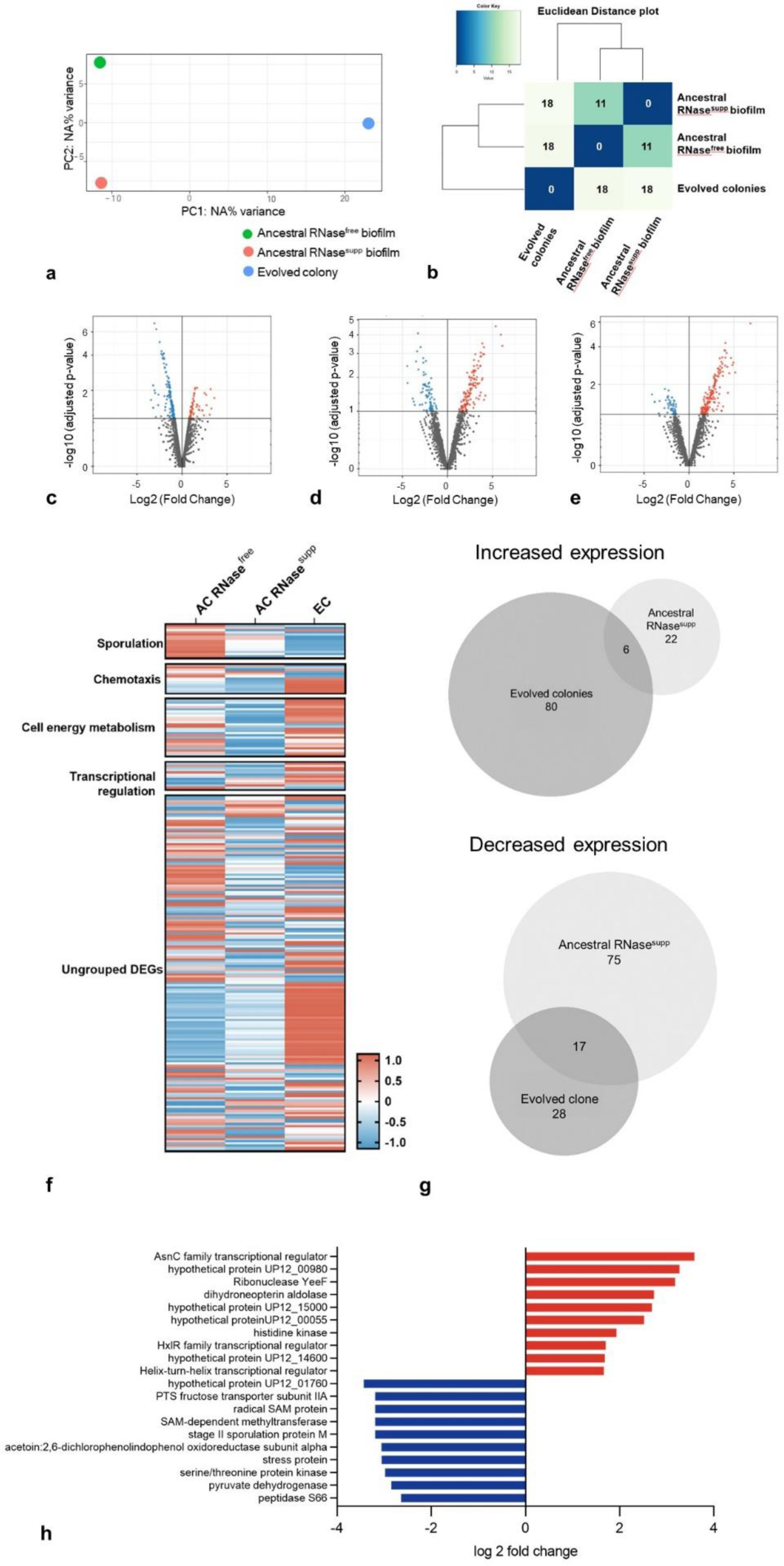
Effect on gene expression of *B. pumilus* cultivation in the presence of RNase. Analysis of differentially expressed genes (DEGs) (fold change > 2; p < 0.05) of cells from ancestral RNase^supp^ biofilms and evolved colonies when compared with those of ancestral RNase^free^ biofilms. (A) PCA plot, (B) Euclidian distance, (C-E) Volcano plots of all differentially expressed genes between (C) ancestral RNase^free^ biofilm and ancestral RNase^supp^ biofilm, (D) ancestral RNase^free^ biofilm and evolved clone, (E) ancestral RNase^supp^ biofilm and evolved clone, (F) gene expression heatmap of positively and negatively regulated DEGs, (G) Venn diagram showing the number of DEGs with increased or decreased expression of ancestral RNase^supp^ biofilm when compared to evolved colonies, (H) top 10 upregulated and downregulated genes when compared ancestral RNase^free^ biofilm vs ancestral RNase^supp^ biofilm.

Next, volcano plots of the log2 (fold change) versus –log10 (p value) show that many of the differentially upregulated and downregulated changes were highly significant in ancestral RNase^supp^ biofilms and evolved colonies when compared to ancestral RNase^free^ biofilms and between each other (Fig. 5 C,D,E). When comparing 120 differentially expressed genes (DEGs) between ancestral RNase^free^ and ancestral RNase^supp^ biofilms, we found that 28 were upregulated and 92 were downregulated. Meanwhile, 132 genes were differentially (fold change > 2) expressed between ancestral RNase^free^ biofilms and evolved colonies, with 86 upregulated and 45 downregulated DEGs (Fig. 5F). The data clearly show differential clustering of gene expression between ancestral RNase^free^ biofilms, ancestral RNase^supp^ biofilms and evolved colonies. Ancestral RNase^supp^ biofilms were downregulated for sporulation, chemotaxis, cell energy metabolism, and transcriptional regulation, compared with to ancestral RNase^free^ biofilm. When compared to both ancestral probes, evolved colonies had more underexpressed genes related to sporulation, while chemotaxis, cell energy metabolism, and transcriptional regulation were overexpressed. (Fig. 5F; Supplementary Table 1). Notably, among chemotaxis-related proteins, evolved colonies showed upregulation of BPUM_2024 pilus assembly protein PilZ with cyclic-di-GMP-binding activity (^36^).

We observed that evolved colonies displayed upregulation of numerous bacteriophage proteins (Supplementary Table 1). Additionally, the number of hypothetical proteins whose expression was significantly altered following the cultivation of bacteria on RNase-supplemented agar was increased. Thus, among 120 DEGs of ancestral RNase^free^ biofilms, 34 proteins were hypothetical (28.3%), and among 132 DEGs of evolved colonies, 36 proteins (27.5%) were hypothetical that is almost three times more enriched than the mean number of hypothetical proteins in the *B. pumilus* proteome, in which only 339 out of 3,608 proteins (9.3%) are classified as hypothetical.

(https://www.ncbi.nlm.nih.gov/genome/browse/#!/proteins/440/368487%7CBacillus%20pumilus/Un/hypothetical). The numbers of unique and overlapping DEGs revealed that only a small fraction of upregulated DEGs were shared between ancestral RNase^supp^ biofilms and evolved colonies when compared to ancestral RNase^free^ biofilms (Fig. 5G).

Next, we analyzed the top 10 genes that were significantly differentially expressed in ancestral RNase^supp^ biofilms when compared to ancestral RNase^free^ biofilms (Fig. 5H, Supplementary Table 1). We found that a number of transcriptional regulators (asnC, HxlR, FZC68_10120) were upregulated, along with the ribonuclease YeeF and genes related to cell division (walk, FolB). We also found downregulated genes that catalyze methylations (queA, BPUM_1881), sporulation (SpoIIM), and acetyl coenzyme A and pyruvate metabolism (acoA_2, BPUM_0452).

To explore the transcriptomic alterations that underlie directed migration of evolved colonies, we next compared the gene expression of ancestral RNase^supp^ biofilms to those of evolved colonies. RNA-seq analysis revealed 121 total DEGs, of which 101 were upregulated and 20 were downregulated in evolved colonies when compared to ancestral RNase^supp^ biofilms (Supplementary Table 1).

Next, we aimed to decipher the roles of the DEGs in the directional migration of evolved colonies. In support of the observed biofilm directional migration, we found that 19 genes associated with chemotactic signaling pathways control were upregulated in evolved colonies when compared to ancestral RNase^supp^ biofilms (Fig. 6A). Among these, we found genes encoding proteins involved in swarming motility, the flagellar system, and chemotaxis, including methyl-accepting chemotaxis proteins (MCPs) (^37^). Moreover, expression of phosphotransferase (PTS) transporters that represent sensory circuits to different carbohydrates through McpB and McpC chemoreceptors was also increased (^38^).

**Figure 6.**
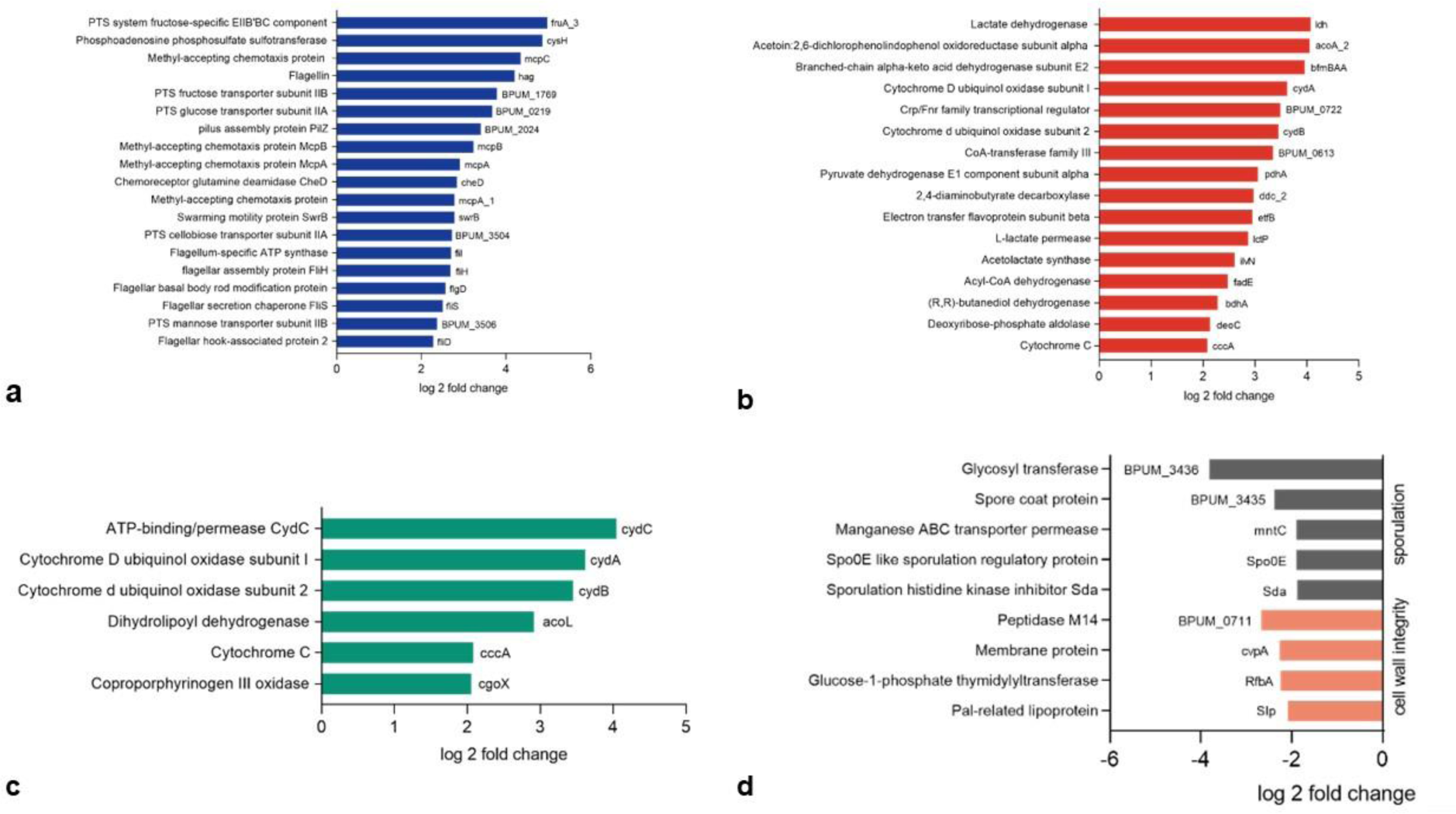
Differentially expressed genes of evolved colonies when compared to ancestral RNase^supp^ biofilms. (A) Genes related to chemotaxis. (B) Genes related to anaerobic metabolism. (C) Genes related to cell redox homeostasis. (D) The top 10 downregulated genes.

Along with these alterations in evolved colonies when compared to ancestral RNase^supp^ biofilms we observed upregulation of many other DEGs. Thus, the expression of critical anaerobic transcriptional units associated with lactate and pyruvate metabolism, which are known to be used as carbon and an energy sources to support bacterial growth were increased (Fig. 6B) (^39^). Also, evolved colonies were characterized by an upregulation of genes implicated in anaerobic respiration, including those associated with lactate formation and formation of 2,3-butanediol via acetoin on the lctEP locus (ldh, lctP) and alsSD operon (ilvN, ddc_2) (^40,41^). Moreover, acoA_2, which catalyzes cleavage of acetoin into acetate and acetaldehyde, was also overexpressed^42^. We also found a significant increase in the expression of numerous other genes related to anaerobic energy metabolism, including those involved in the conversion of branched amino acids and pyruvate to acetyl-CoA (pdhA, bfmBAA), as well as others which enable *Bacillus* spp. to utilize nitrate as an alternative electron acceptor (fade, BPUM_0613, etfB, dhA and BPUM_0722 (^40,43–46^). We discovered upregulation of other genes related to energy metabolism (deoC, cydA, cydB, cccA) that participate in ATP production under limited oxygen conditions (^47–51^). Interestingly, they may play a dual role, as being viewed as oxidoreductive enzymes in evolved colonies and along with overexpressed cydC, acoL, and cgoX, they may maintain redox homeostasis under the activation of anoxic respiration (Fig. 6C) (^47,50,52^). Also, in evolved colonies we observed increased expression of the RecT recombinase, which regulates chromosomal insertions, (Supplementary Table 1) (^53^).

Among the 20 genes for which expression was downregulated in evolved colonies, five of these genes are associated with sporulation, including sda, spo0E, BPUM_3435, mntC, and BPUM_3436 (Fig. 6D) (^54–56^). Other underexpressed genes included those which have previously been shown to be related to cell wall integrity.

### Differences in biochemical activity between ancestral RNase^free^ biofilms, ancestral RNase^supp^ biofilms, and evolved colonies

The biochemical profile of bacteria from ancestral RNase^free^ biofilm, ancestral RNase^supp^ biofilm and evolved colonies were analyzed with an automated system VITEK® 2 (Table 2). We found that ancestral RNase^supp^ biofilms were positive for leucine arylamidase, β-D-glucosidase, and D-glucose in contrast to ancestral RNase^free^ biofilm.

When comparing bacteria from evolved colonies and ancestral RNase^supp^ biofilms, we observed dysregulation of several enzymes, including activation in evolved colonies of urease, tyrosine arylamidase, and arginine hydrolase, which are part of basic energy metabolism and stress resistance (^57–60^). Cells from evolved colonies differed from those from both ancestral RNase^supp^ biofilms and ancestral RNase^free^ biofilms in that they did not ferment β-n-acetylglucosaminidase, D-maltose, Beta-galactosidase, or L-arabinose. However, some characteristics of evolved colonies (that were also cultivated on RNase-supplemented agar) differed from ancestral RNase^supp^ biofilms but exhibited the same utilization pattern as ancestral RNase^free^ biofilms, and were negative for bacterial energy metabolism enzymes D-glucose and β-D-glucosidase (^61,62^).

### Identification of extracellular DNA as a chemoattractant

To determine the factor within human blood that acts as a chemoattractant for directional migration in our model, we separately compared plasma, whole blood, and blood cells. We identified that directional migration occurred only toward plasma (Fig. 7A) and whole blood (Fig. 7B), but not toward blood cells (Fig. 7C).

**Figure 7:**
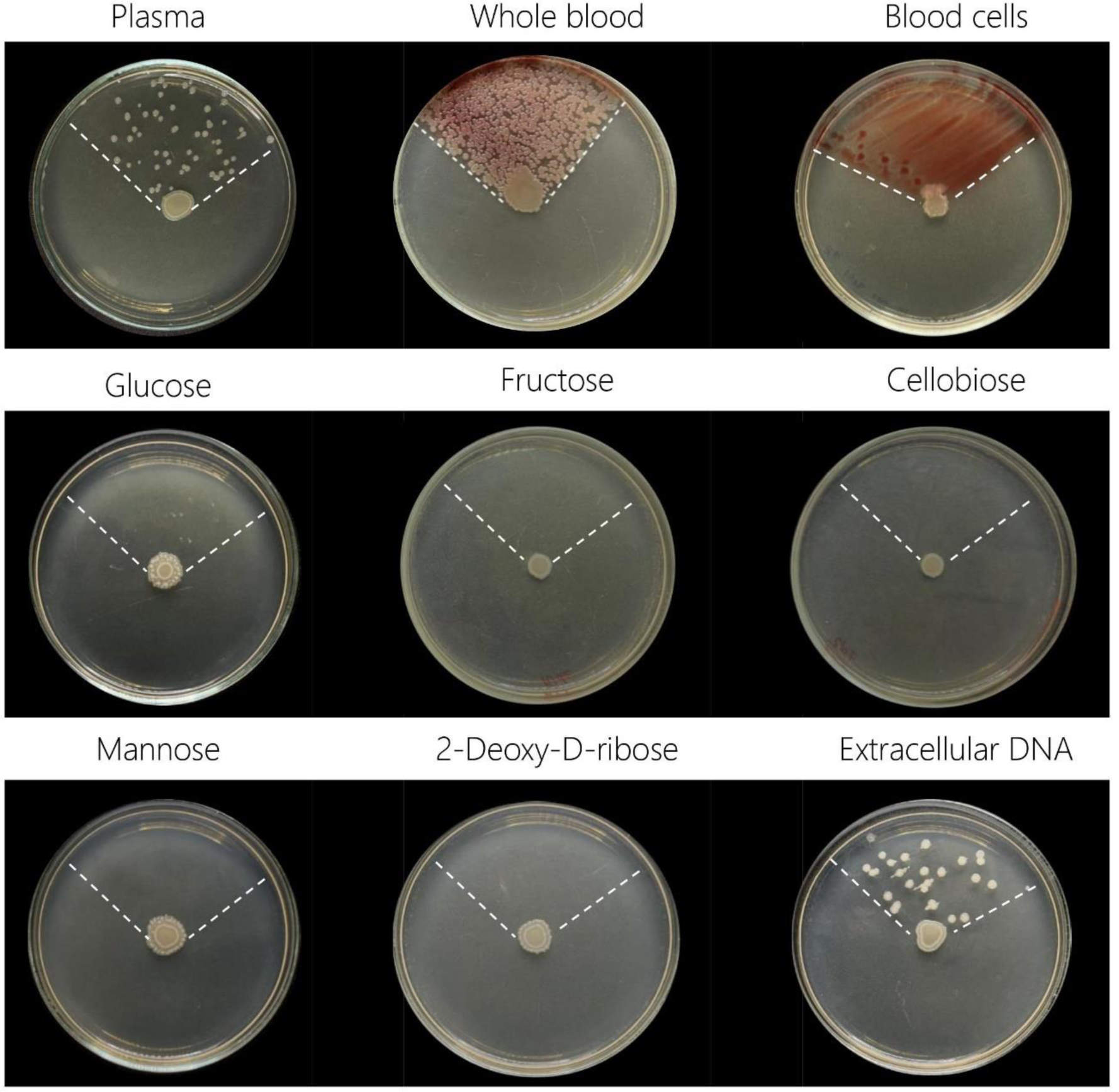
Comparison of *B. pumilus* directional migration in response to different potential chemoattractants cultivated on RNase-supplemented agar. The dotted area denotes the application area for potential chemoattractants. Pictures are from representative experiments, which were performed in triplicate.

Since transcriptomic data revealed the upregulation of PTS and McpC genes involved in sensing different sugars to activate chemotaxis, we next studied whether plasma carbohydrates could be chemoattractants that trigger biofilm dispersal and resettlement. We chose to test glucose, fructose, cellobiose, and mannose as chemoattractants because PTS genes involved in their recognition were upregulated, but none of them triggered directional migration of *B. pumilus* (Fig. 7D,E,F,G). Next, we wondered if the DNA in the plasma could be a chemoattractant that triggers biofilm directional migration given that it has a sugar backbone made of deoxyribose. While the addition of 2-Deoxy-D-ribose did not trigger directional migration, the addition of blood-derived extracellular DNA promoted directional migration of *B. pumilus* toward it (Fig. 7H,I). Collectively, point out that in these experiments, plasma DNA acted as a chemoattractant for *B. pumilus* in the experiments. Importantly, the addition of extracellular DNA to ancestral RNase^free^ biofilms did not affect biofilm dispersal (Supplementary Figure 4).

## DISCUSSION

After our recent demonstration of the role of extracellular RNA as the component of the Universal Receptive System in bacterial migration and biofilm dispersal, the detailed mechanism of such a regulation remained enigmatic (^28^). In the present study, we revealed in detail how the destruction or inactivation of extracellular RNA in the presence of a chemoattractant can activate chemotaxis and directional migration of *B. pumilus*, including dispersal and resettlement of developing and mature biofilms. This is particularly unusual because, while dispersal is typical for mature biofilms when elevated cell density causes migration toward external stimuli, we show in this study that loss of extracellular RNA overrides normal biofilm regulation, making cells from biofilms of any age ready to migrate (^63,64^). What also makes the results unexpected is that the bacteria migrated on a solid 1.5% agar, whereas normally, bacteria lack the ability to migrate long distances on such a dense medium (^65^). We dissected this process by the hour and found that bacteria on agar supplemented with RNase and added chemoattractant got the signal to migrate within the first 0.5 h after plating to the agar. This means that within this time frame, bacteria initially sensed the loss of extracellular RNA and then switched on flagella-driven chemotaxis and cellular decision-making, which are necessary for migration. This resulted in the formation of some subsets of migrating “runner” cells in what we call a “panic state” that formed evolved colonies.

Based on this observation that the response of bacteria following the modulation of the Universal Receptive System and inactivation of extracellular RNA occur rapidly; we performed transcriptomics (^66^).

he inactivation of extracellular RNA resulted in multiple alterations to bacterial gene expression, with the most profound changes observed in evolved colonies. We observed the upregulation of the genes in evolved colonies that been recognized to influence chemotaxis and the motility of migrating cells, including flagellar-specific genes and the well-known chemotaxis-specific transmembrane proteins McpA, McpB, and McpC (^67^). Notably, among the transmembrane Che-family of chemoreceptors, only the expression of CheD was increased. CheD, which is classified as a chemoreceptor glutamine deamidase, appears to have two separate roles in the chemotaxis of *Bacillus spp.* One of them is realized by deamidation of Mcp-family proteins, and another by affecting the CheC/CheD/CheY adaptation system that is necessary for chemotaxis-related mobility (^68–70^).

It has not escaped our attention that the upregulated pilus assembly protein PilZ possesses c-di-GMP-binding activity. Previous studies have reported that overexpression of c-di-GMP-binding proteins results in a decreased cellular level of c-di-GMP that inhibits flagellar rotation by curbing flagellar motor output (^71–74^). We speculate that PilZ could stimulate motility of the evolved colonies in this study through the c-di-GMP pathway (^75^).

Also overexpressed in evolved colonies were PTS transporters of glucose, fructose, cellobiose, and mannose that act as co-sensors for bacterial signal transduction systems, regulating chemoreception to carbohydrates by transmitting the stimuli generated by PTS substrates to the cytosolic fragment of the McpC (^38,76,77^). Therefore, even though bacteria were cultured on a rich multi-purpose medium with free access to carbon we decided to check if the presence of these sugars could trigger directional migration. As expected, none of these sugars triggered the formation of the evolved colonies, meaning that plasma carbohydrates were not the triggering factors for chemotaxis.

By excluding blood cell elements as potential chemoattractants, we hypothesized that *B. pumilus* in the case of extracellular RNA destruction dispersed towards blood-derived extracellular DNA. We observed that the addition of extracted extracellular plasma DNA also triggered chemotactic-induced *B. pumilus* migration. Moreover, that occurred only if DNA was polymeric—the DNA-related carbohydrate 2-Deoxy-D-ribose did not act as a chemoattractant. Since expression of the RecT recombinase, which regulates chromosomal insertions, was also increased in evolving colonies, we can speculate that polymeric extracellular DNA molecules may be at some point used by the cells for the natural transformation (^53^). These data are in line with our previous discovery that novel TezR receptors formed by extracellular RNA also control genetic alterations, along with orchestrating the interaction of cells with their environment (^28,33^).

Furthermore, we found that although cultured aerobically, evolved colonies showed upregulated expression of multiple genes which are known to help cells adapt to anoxic conditions (^40^). Several groups have reported that *Bacillus* spp. in the absence of oxygen can utilize nitrate as an alternative electron acceptor, with lctEP, alsSD, and pta loci on the chromosome involved in this process. In our study, the ldh and lctP genes from the lctEP locus and the ilvN and ddc_2 genes from the alsSD locus were upregulated (^78–80^). Although there was no alteration in phosphotransacetylase expression encoded by *pta*, which is required for acetate synthesis from acetyl coenzyme A, we found that acoA_2, which performs the same function, was significantly overexpressed (^42^).

Moreover, we observed overexpression of multiple key genes that regulate anaerobic energy metabolism through the acetate and acetyl-CoA pathways (^81^). Although the activation of anaerobic-related energetic pathways under aerobic conditions remains poorly understood, it should provide a significant energetic boost to evolved colonies for migration (^82^). It remains, therefore, an outstanding question about the underlying regulatory pathway since the activation of these genes does not seem necessary and was not anticipated in unchanged gas, environmental, and nutrition conditions. Expectedly, the number of ATP-binding proteins required to release the energy was also overexpressed (^83^).

Biochemical profiles of *Bacillus* from evolved colonies, confirmed that unlike those from ancestral biofilms, they had a positive urease reaction.

Urease is an enzyme that enables *Bacillus* spp. to utilize nitrogen as an energy source, and arginine hydrolase as another reservoir of nitrogen (^57,60,84,85^). Notably, the *B. pumilus* genome does not have a urease gene, so the fact that they were capable of urea hydrolysis indicates that evolved colonies experienced significant genomic alterations (NCBI GenBank assembly accession: GCA_003020795.1).

It is not surprising that such intensive alterations to energetic metabolism resulted in the alteration of genes that catalyze the reduction of reactive substrates. Thus, oxidoreductive enzymes involved in redox homeostasis under anaerobic respiration, including cytochromes, were upregulated in evolved colonies (^50,52,86^). Biochemical analyses confirmed a positive tyrosine arylamidase reaction, which releases tyrosine residues to help cells withstand different stresses (^52^).

Cultivation of bacteria on RNase-supplemented agar resulted in the formation of smaller cells with a higher sporulation frequency in ancestral RNase^supp^ biofilms and evolved colonies. Two things attracted our attention. The first why the level of the sporulation frequency, which was increased across ancestral RNase^supp^ biofilms and evolved colonies, was directly correlated with the downregulation of known sporulation genes. One explanation is that genes were downregulated since the sporulation process was completed; however, this requires additional observation. The second outstanding question is how the evolved colonies exhibited increased sporulation along with increased motility, since this contradicts previous reports that sporulation downregulates expression of motility-related genes (^87^). The DEGs of both ancestral RNase^supp^ biofilms and evolved colonies were over three-fold enriched with hypothetical genes, meaning that the destruction of extracellular RNA triggered an unusual cell response involving genes of unknown functions. Also, we found that bacteriophage genes were upregulated in the evolved colonies; however, no signs of lysogenic bacteriophage infection were observed. Although this finding is in line with data from previous research showing that some cryptic-prophage-encoded products can regulate bacterial hosts’ metabolism and division, the reason for such profound bacteriophage activation is unclear (^88–90^).

Future research should aim to improve our understanding of the role of these hypothetical genes in the regulation of cell migration, and of the molecular pathways between the loss of extracellular RNA and cell response, and how these signals are transmitted. Taken together, this study adds another line of evidence that extracellular RNA as a tool of New Biology in bacterial biofilms is a unique and functionally active part of the extrabiome. It highlights the previously unexplored role of the Universal Receptive System in governing the responses of cells and microbial communities to the outer environment (^91^).

Our results suggest that the extracellular RNA of the Universal Receptive System can be used to modulate various cell responses related to directional bacterial migration and provide additional approaches for regulating biofilm dispersal.

## MATERIALS AND METHODS

### Reagents

Bovine pancreatic DNase I (2,200 Kunitz units/mg), RNase A, glucose, fructose, cellobiose, mannose, and 2-Deoxy-D-Ribose, were all obtained from Sigma-Aldrich (St. Louis, Missouri, USA). Blood specimens were collected in K2-EDTA blood collection tubes from volunteers after giving written consent, and the probes were centrifuged to isolate plasma and RBCs. The study was approved by the institutional review board of the Human Microbiology Institute (# VB-022376).

### Bacterial strains and culture conditions

For biofilm formation, overnight culture of *Bacillus pumilus* VT1200 (provided by Dr. V. Tetz) in LB broth (Oxoid, Hampshire, UK; Sigma-Aldrich) were harvested by centrifugation at 4,000 rpm for 15 min (Microfuge 20R; Beckman Coulter, La Brea, California, USA), supernatant was discharged and the pellet was washed twice and resuspended in phosphate-buffered saline (PBS, pH 7.2) (Sigma-Aldrich). Then, 25 µL of bacterial suspension containing 5.5 log10 cells was inoculated into the center of 90-mm glass Petri dishes filled with 1.5% TGV agar (Human Microbiology Institute, New York City, New York, USA) and cultivated aerobically at 37 °C.

To study the destruction of extracellular RNA, after autoclaving at 121 °C for 20 min, the agar was cooled down to 45 °C. RNase A at a final concentration 50.0 µg/mL, or anti-RNA antibodies, were added, mixed, and poured onto the surface of the cooled agar.

### Directional migration and biofilm dispersal

To study directional dispersal, we added 50 µL of chemoattractant—whole blood, plasma, red blood cells (RBCs), glucose (10 mM), fructose (10 mM), cellobiose (10 mM), mannose (10 mM), 2-Deoxy-D-Ribose (10 mM), or 100 ng of total DNA isolated from plasma—to a sector comprising 1/4-1/3 of the plate using a sterile inoculation loop or a cell spreader (Cole Parmer, USA). In some experiments, 50 µL of plasma was added to a 24-cm-long glass plate in the pattern of a pathway. This pathway was designed to guide the growth of bacteria from the original inoculum along the chemoattractant pathway within the corridor created by plastic inserts placed in the agar. Subsequently,.

an aliquot containing 5.5 log10 *B. pumilus* VT1200 in 25 µL was placed in the center of the 90-mm plates or 24-cm-long glass plates, which contained TGV agar with the chemoattractant already added and dried. The aliquot of *B. pumilus* VT1200 was added as a drop and left to dry, ensuring that it did not disperse.. The plates were incubated at 37 °C for 24 h.

To study the directional migration or preformed biofilms, 24-h-old biofilms were grown on the agar surface as described above, without the addition of a chemoattractant. Next, chemoattractant was added a sector comprising 1/4 of the plate, and the plates were cultivated at 37 °C for an additional 24 h.

### Directional migration in four-compartment Petri dishes

Two Petri dishes were filled with TGV agar (Human Microbiology Institute, New York City, New York, USA). In the first dish, agar was supplemented with RNase A dissolved in 0.9% NaCl aqueous solution at a final concentration 50.0 µg/mL. The agar in the second dish was left untreated. In both Petri dishes, the agar was cut into four equal pieces and placed onto an empty Petri dish, resulting in two pieces of RNase-supplemented agar and two pieces of agar without supplementation. Next, a 6-mm well was cut from the center of the agar and removed. Fresh unsolidified TGV agar at 60 °C was used to fill the central well and gaps between agar pieces. Subsequently, four agar sections were separated accordingly with plastic impenetrable partitions.

The agar without RNase was supplemented with 250 µL of 0.9% NaCl aqueous solution, which was distributed within a ¼ section of the Petri dish with cell spreader (Cole Parmer, USA). On two sections of the agar (one supplemented with RNase A and one without), 250 µL of plasma from health volunteers was added and distributed with a cell spreader (Cole Parmer, USA). Thus, the experimental four-compartment plate contained the following sections: (1) 0.9% NaCl aqueous solution, (2) plasma (3) RNase 50.0 µg/mL, or (4) plasma and RNase 50.0 µg/mL.

After drying, an aliquot containing 5.5 log10 *B. pumilus* VT1200 in 25 µL was placed on the center of the agar and the plates were cultivated at 37 °C for 24 h.

### Light microscopy

Cells were imaged using an Axioscope plus microscope (Carl Zeiss, Jena, Germany) under a 100 × oil objective (ApoPlan 100× 1.25 oil Ph3 [Zeiss]) using a Canon 6 camera and Fiji/ImageJ software^92^. Cell size was determined following Gram staining (Sigma-Aldrich). Values were expressed in pX2. The average number of spores was estimated by counting spores in over 20 fields of view.

### Camera and software

A video of bacterial migration on a 24-cm-long plate, prepared as described above, was captured using a Canon 6 camera set for video recording inside an incubator at 37°C. To minimize the file size, the low-resolution setting was chosen. The captured videos covering the first 18 hours of bacterial dispersal were converted to a movie format and accelerated to 64x speed using the open-source video transcoder Online Video Cutter.

### Colony characteristics assay

*B. pumilus* VT-1200 colonies were photographed using a Leica MZ9.5 stereomicroscope (Leica Biosystems, Newcastle, UK). The area of colonies was measured using Fiji/ImageJ software and expressed in pX2. The number of CFUs per 24-h-old colonies was measured using the serial dilution plating method, as previously described. The optical density of 24-h-old colonies was determined in triplicate using a spectrophotometer at 600 nm (SmartSpec Plus, Bio-Rad, USA).

### DNA extraction

Before nucleic acid extraction, plasma was filtered through a 0.22-μm filter (Millipore Corp., Bedford, Massachusetts, USA). Extracellular DNA from plasma was extracted using a NucleoMag® DNA Plasma Kit (Thermo Fisher Scientific Inc., Waltham, Massachusetts, USA) according to the manufacturer’s instructions.

### Generation of anti-RNA antibodies

Anti-RNA antibodies were obtained as previously described by our group (^28,33^). Briefly, to isolate RNA, the supernatant of 24 h *B. pumilus* biofilms was filtered through a 0.22-μm filter (Millipore Corp., Bedford, MA, USA) and the extracellular RNA was extracted by using an RNeasy Mini Kit (Qiagen, Valencia, CA) according to the manufacturer’s instructions. Antibodies against extracellular RNA were obtained after immunization of 4-month-old New Zealand White rabbits with RNA and complete Freund’s adjuvant, according to the Cold Spring Harbor protocol for standard immunization of rabbits (^93^).

### Total RNA extraction, purification, Illumina library preparation, and sequencing

Following 12 h of growth on RNase-free or RNase-supplemented agar, ancestral and evolved biofilms were collected and gently washed twice with PBS. Cells were harvested at 4000 × *g* for 15 min (Microfuge 20R, Beckman Coulter) and resuspended in PBS. RNA was extracted from bacteria using an RNeasy Mini Kit (Qiagen) according to the manufacturer’s instructions. RNA quality was assessed spectrophotometrically at 230/260/280 nm with the NanoDrop OneC spectrophotometer (ThermoFisher Scientific).

Ribodepletion was performed using the Ribo-Zero Magnetic Gold kit (Epicenter, Madison, Wisconsin, USA) according to the manufacturer’s guidelines. Transcriptome sequencing libraries were prepared from RNA using the TruSeq Stranded Total RNA Library Prep Kit according to the manufacturer’s instructions, and sequenced using a 2 × 150 nucleotide paired-end strategy (Illumina NextSeq 500, Illumina, San Diego, California, USA).

### RNA-seq data processing

All of the reads from the Sequencing experiment were mapped to the reference genome (B. pumilus 8325-4)using the Bowtie2 (v2.2.4) (PMID: 22388286) and duplicate reads were removed using Picard tools (v.1.126) (http://broadinstitute.github.io/picard/). Low quality mapped reads (MQ<20) were removed from the analysis. The read per million (RPM) normalized BigWig files were generated using BEDTools (v.2.17.0) (PMID: 20110278) and the bedGraphToBigWig tool (v.4). The read count tables were generated using HTSeq (v0.6.0) (Anders et al., 2015), normalized based on their library size factors using DEseq2 (Love et al. 2014), and differential expression analysis was performed. Transcripts with an adjusted p value of < 0.05 and log2 fold change value of ± 1.0 (2 fold) were considered significantly differentially expressed. To compare the level of similarity among the samples and their replicates, we used two methods: principal-component analysis and Euclidean distance-based sample clustering. All the downstream statistical analyses and generating plots were performed in R environment (v3.1.1) (https://www.r-project.org/).

### Biochemical analyses

Biochemical analyses were performed using the colorimetric VITEK® 2 COMPACT 30 system (BioMérieux, Marcy l’Étoile, France) with BCL (spore-forming bacilli) Data were analyzed using VITEK® 2 software version 7.01.

### Statistics

At least three biological replicates were performed for each experimental condition unless stated otherwise. Each data point was denoted by the mean value ± standard deviation (SD). A two-tailed t test was performed for pairwise comparisons and p ≤ 0.05 was considered significant. Bacterial quantification data were log10-transformed prior to analysis. Statistical analyses for the biofilm assays and hemolysin tests were performed using Student’s t test. GraphPad Prism version 9 (GraphPad Software, San Diego, California, USA) or Excel 10 (Microsoft, Redmond, Washington, USA) were applied for statistical analysis and illustration.

## ACKNOWLEDGEMENTS

We would like to thank the Genome Technology Center (GTC) for expert library preparation and sequencing, and the Applied Bioinformatics Laboratories (ABL) for providing bioinformatics support and helping with the analysis and interpretation of the data. GTC and ABL are shared resources partially supported by the Cancer Center Support Grant P30CA016087 at the Laura and Isaac Perlmutter Cancer Center. This work has used computing resources at the NYU School of Medicine High Performance Computing (HPC) Facility.

## Funding Sources

NCI/NIH Cancer Center Support Grant P30CA016087

**Supplementary Figure 1:** The effect of extracellular RNA and DNA destruction on biofilm migration toward plasma used as chemoattractant on 1.5% agar.

**Supplementary Figure 2:** Microscope image of bacteria from 24-h-old ancestral RNase^free^ biofilms, ancestral RNase^supp^ biofilms, and evolved colonies. The microscopy of cells from (A) ancestral RNase^free^ biofilm (B) ancestral RNase^supp^ biofilm (C) evolved colonies at 2 cm from inoculum, (D) evolved colonies at 4 cm from inoculum, (E) evolved colonies at 6-cm from inoculum. The dotted area is the plasma application area (bars=10µm).

**Supplementary Figure 3:** The effect of extracellular RNA on biofilm migration toward plasma used as chemoattractant on 1.5% agar. (A) Characteristics of bacteria from 12-h-old ancestral RNase^free^ biofilms, ancestral RNase^supp^ biofilms, and evolved colonies. (B-E) Microscope image of bacteria from 12-h-old ancestral RNase^free^ biofilms, ancestral RNase^supp^ biofilms, and evolved colonies. The microscopy of cells from (B) ancestral RNase^free^ biofilm (C) ancestral RNase^supp^ biofilm (D) evolved colonies at 2 cm from inoculum, (E) evolved colonies at 4 cm from inoculum. The dotted area is the plasma application area (bars=10µm). (F) Characteristics of bacteria from 12-h-old ancestral RNase^free^ biofilms, ancestral RNase^supp^ biofilms, and evolved colonies.

**Supplementary Figure 4:** Effect of plasma DNA as a chemoattractant added to ancestral RNase^free^ biofilms.

**Supplementary Data 1:** List of differentially expressed genes of ancestral RNase^free^ biofilms, ancestral RNase^supp^ biofilms, and evolved colonies.

**Supplementary Video 1:** Video image of *B. pumilus* migration. The video shows a time-lapse (64x speed) of 12 hours of *B. pumilus* migration, with plasma used as a chemoattractant added to the agar surface in the form of a path to direct bacterial chemotaxis.

## Tables

**Table 1:** Enzymes with altered activity between bacteria from ancestral RNase^free^ biofilms, ancestral RNase^supp^ biofilms, and evolved colonies.

| Enzymes<br>activated across probes<br>differently | Ancestral<br>RNase<sup>free</sup><br>biofilm | Ancestral<br>RNase<sup>supp</sup><br>biofilm | Evolved<br>clone |
| --- | --- | --- | --- |
| Leucine arylamidase | - | + | + |
| Tyrosine arylamidase | - | - | + |
| D-glucose | - | + | - |
| β-n-acetylglucosaminidase | + | + | - |
| D-maltose | + | + | - |
| Urease | - | - | + |
| Beta-galactosidase | + | + | - |
| Arginine hydrolase 1 | - | - | + |
| Beta-d-glucosidase | - | + | - |
| L-arabinose | + | + | - |

Biochemical characteristics were obtained by VITEK2 COMPACT with the GP ID card; +: positive;-: negative reaction; for congruent results, see text.

