## Supplementary figures and images for "Previously unknown regulatory role of extracellular RNA on bacterial directional migration"

### Supplementary Figure 1

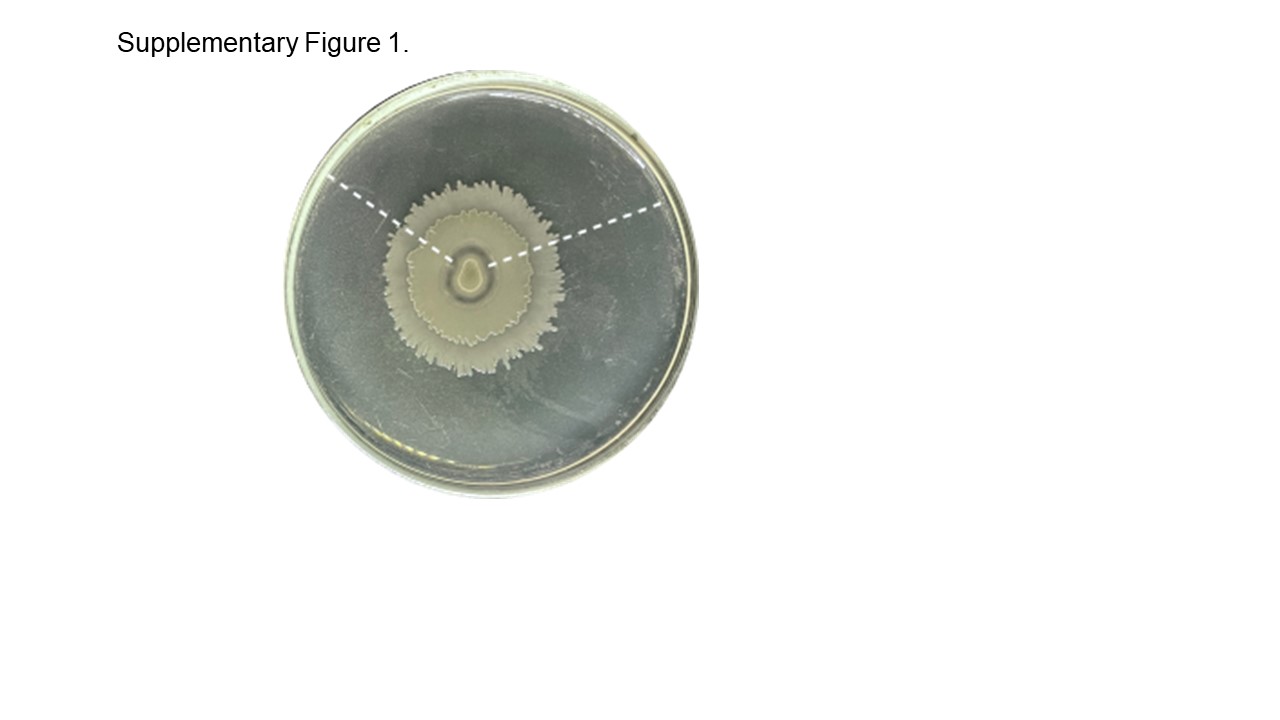

### Supplementary Figure 2

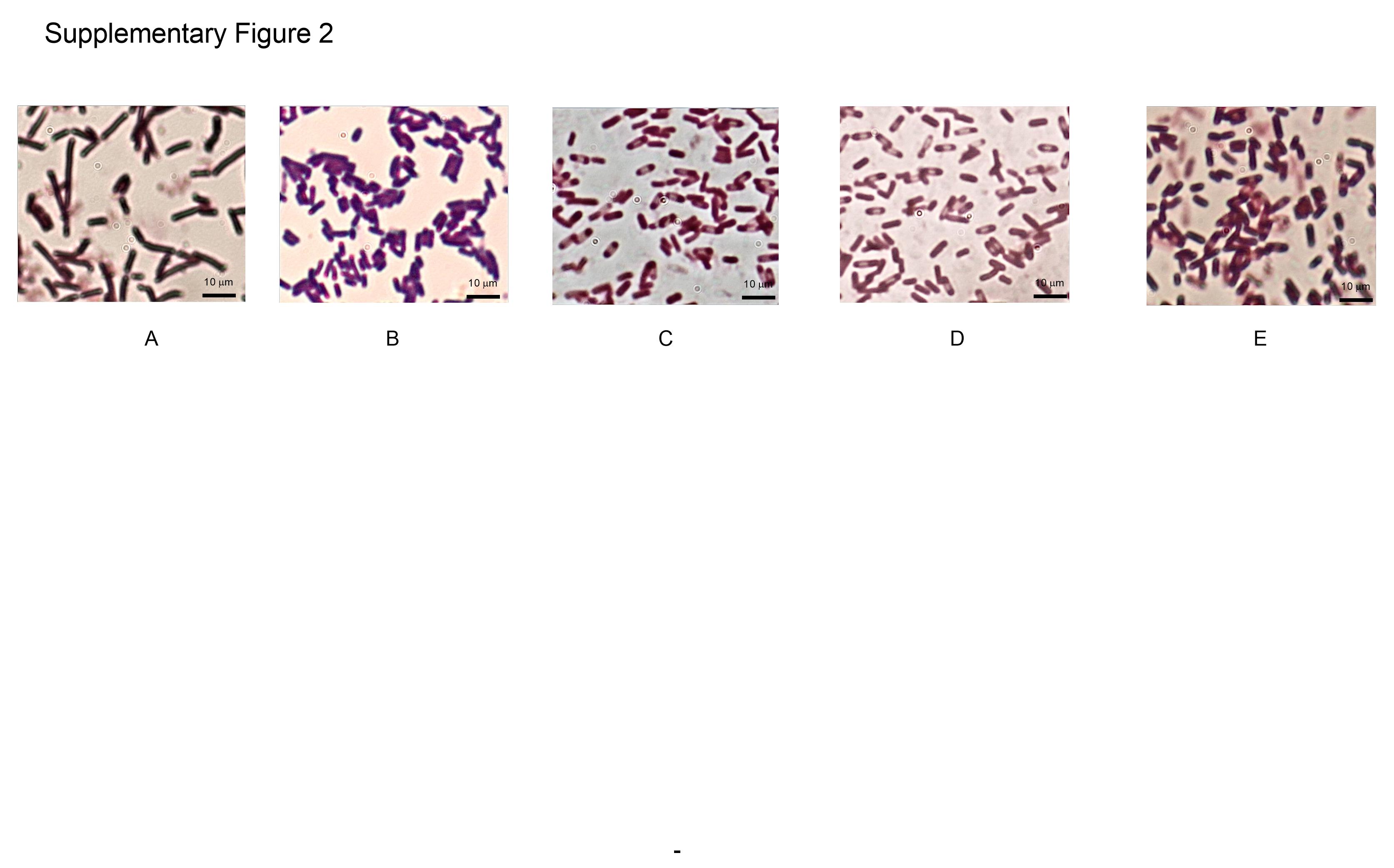

### Supplementary Figure 3

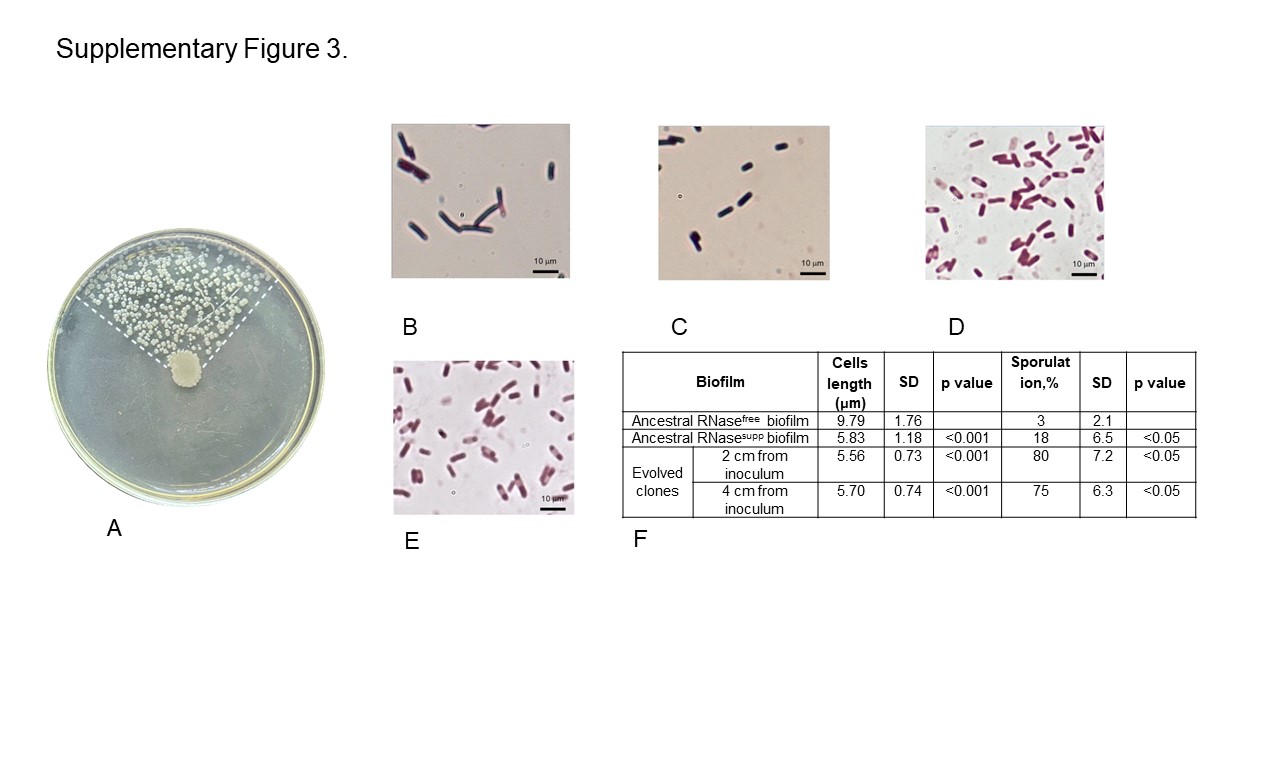

### Supplementary Figure 4

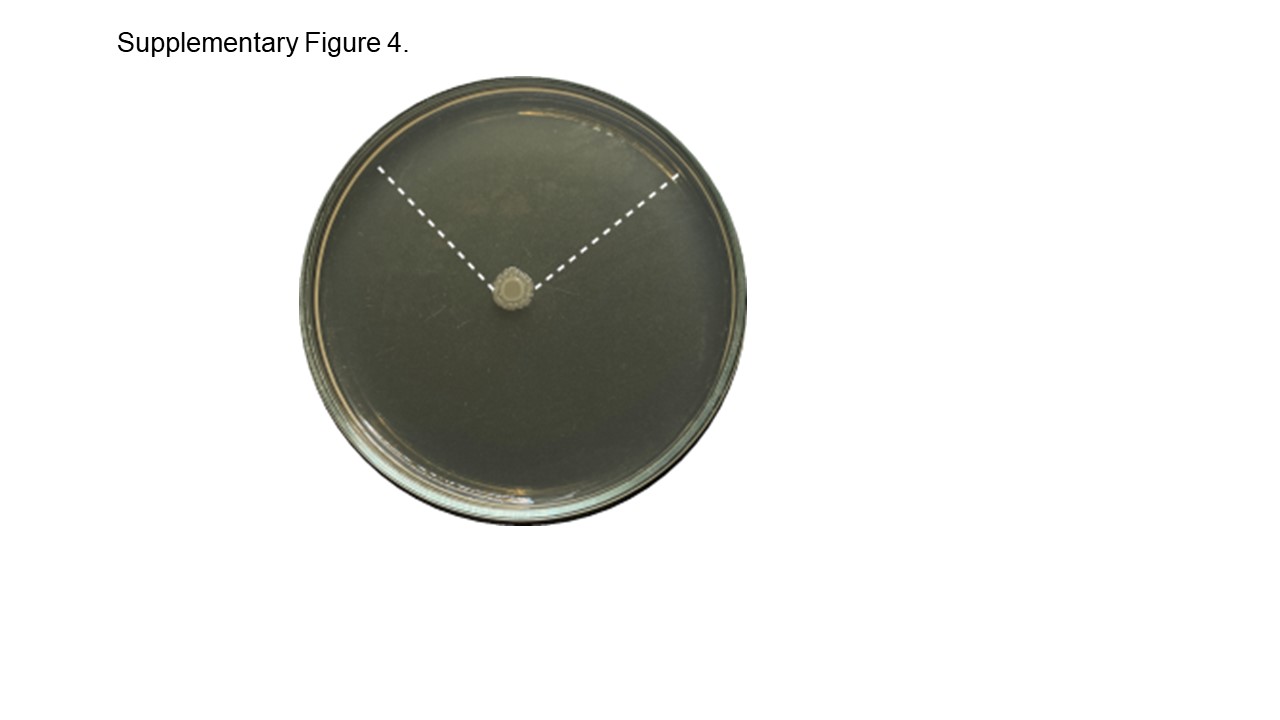
